# Interplay between m6A modification and overall transcripts quantity: Impacts on mRNA composition in plant stress granules

**DOI:** 10.1101/2023.12.14.569339

**Authors:** Dawid Jakub Kubiak, Michal Wojciech Szczesniak, Karolina Ostrowska, Dawid Bielewicz, Susheel Sagar Bhat, Katarzyna Niedojadlo, Zofia Szweykowska-Kulinska, Artur Jarmolowski, Rupert George Fray, Janusz Niedojadlo

## Abstract

Stress granules (SGs) are cytoplasmic structures that emerge in response to unfavorable environmental conditions. They contain a rich pool of RNA, including non-translated mRNA. The mechanisms governing transcripts accumulation in SGs is only partially understood. Despite the recognized role of m6A in plant transcriptome regulation, its impact on SGs’ composition and assembly remains elusive. We examined the formation of SGs, the presence of m6A, and the transcription-level-dependent localization of selected mRNAs within these granules during hypoxia in the roots of *Lupinus angustifolius* and *Arabidopsis thaliana*. In lupine, SGs exhibit a distinctive bi-zonal structure, comprising of a ring and a central area with differences in ultrastructure and composition. Following the transcriptome analysis, mRNAs were selected for examination of their localization in SGs and m6A levels. Transcripts from genes responsive to hypoxia (ADH1 and HUP7) exhibited significant lower levels of m6A compared to housekeeping genes but only ADH1 was not present in SGs. HUP7 mRNA with low quantity of m6A, is present both in the SGs and cytoplasm probably due to extremely high expression level. It was also shown that the amount of m6A in SGs was higher than in the cytoplasm only in the first hours of hypoxia and then decreased. In mutants of A. thaliana with reduced level of m6A, formation and quantity of SGs were studied. In this line, ECT2 was not observed and poly(A) RNA levels were slightly reduced in SGs. Additionally the number of SGs was lower than that of the wild type. In summary, our findings demonstrate the limited impact of m6A modification on SGs assembly. However the interplay between m6A modification and the overall transcript quantity in the cytoplasm plays a regulatory role in mRNA partitioning into SGs.

## Introduction

The cell’s response to adverse environmental conditions requires regulation of gene expression. Recently, it has been shown that spatial regulation of this process during stress is important (de Nadal et al., 2011). The structural manifestation of this phenomenon is storage of RNA in stress granules (SGs) in the cytoplasm. The mRNA accumulated in SGs is separated from the translation complexes. This mechanism enables the reduction of protein synthesis and regulates the selectivity of the translation process. The RNA accumulated within these structures can be utilized following end of stress (Kearly et al., 2022).

SGs are dynamic membrane-less cytoplasmic structures that formed in plant and animal cells exposed to abiotic and biotic stress (Maruri-López et al., 2021). The breakdown of polysomes results in the release of mRNA and the formation of mRNA-ribonucleoprotein complexes (mRNPs). It is commonly believed that stress-induced inhibition of translation is considered a key trigger for the formation of stress granules (Sorenson and Bailey-Serres 2014, Hofmann et al., 2021). Previous studies in mammals and yeast suggest that a several-step mechanism, conserved among eukaryotes and driven by liquid-liquid phase separation (LLPS), is responsible for formation and dynamics of SGs. There is limited information on stress granules formation in plants, but several studies provide some evidence supporting mechanism proposed for animal cells (Allen and Strader 2022; Gutierrez-Beltran et al., 2021). The first step in the formation of SGs is nucleation, during which a high concentration of mRNPs and a combination of protein-protein, RNA-RNA and protein-RNA interactions induces LLPS of mRNP complexes lead to assembly of the SGs core (Niewidok et al., 2018). An important feature of SGs core proteins, known in plant and animal models, is their multi-domain architecture, including oligomerization domains (ODs), RNA-binding domains (RBDs) and interactions between proteins (Sanders et al., 2020). In contrast to animals, the effect of oligomerization of SGs core proteins in the nucleation process in plants is poorly studied. During core growth the secondary structure of RNA is of particular importance, and can act as a binding factor to liquid condensates. A correlation between RNA structure and affinity for binding to proteins has been demonstrated (de Groot et al., 2019). However, the transcriptome of SGs and the role of RNA structure in their formation is poorly understood in plants. The final step in SGs formation is shell condensation. Once the core structure is stabilized, additional mRNPs, proteins, small molecules and nucleotides are recruited as shell components, forming microscale SGs. The proteins that make up the shell mostly lack RNA-binding activity, and are dependent on the organism, cell type, and type of stress (Markmiller et al., 2018). In the final step, microscale SGs combine to form a multicore structure immersed in a single shell (Wheeler et al., 2016). The distinction between cores and shells in plants is much less obvious than in animals. Studies on animal cells have shown that the formation and degradation of SGs is an active process, requiring energy in the form of ATP (Shao et al., 2017). In plants presence of ADP and proteins with ATPase activity in SGs has been reported, however, it is unknown whether these proteins are active and what role ATP plays (Kosmacz et al., 2019). Microtubule-dependent transport is important in the process of SGs formation in both plants and animal models. Disruption of cytoskeletal elements by a microtubule-actin polymerization inhibitor results in blocking the fusion step of SGs (Hamada et al., 2018). Despite several similarities with the animal model, our understanding of the process of stress granules formation in plants remains limited, with substantial knowledge primarily confined to *A. thaliana*.

It has been shown that SGs may be a storage site for translation-ready mRNA not associated with ribosomes in stress conditions. Surviving the conditions of abiotic stress is related to the limitation of processes that require large amounts of energy in the cell (Das et al., 2022). Therefore, under abiotic stress conditions, it is necessary to reduce the intensity of translation. Nonetheless, cellular response to stressful conditions necessitates the presence of specific proteins. Currently, the exact mechanisms governing selective translation remain unclear (Merchante et al., 2017). Separation of mRNA molecules from the translational apparatus and their sequestration in stress granules may be a strategy to allow the translation of only selected proteins during stress (Hofmann et al., 2021). Under abiotic stress conditions, the translation initiation factor eIF2a is phosphorylated. This inhibits the translation process and the breakdown of polysomes. mRNAs are released and likely transported to SGs (Hofmann et al., 2021). It has been shown that increasing the stability of polysomes with cycloheximide leads to the disassembly of SGs, while the degradation of polysomes with puromycin promotes SGs formation (Kedersha et al., 2000; Merret et al., 2017). Based on the above observation, it is assumed that transcripts whose translation has been inhibited are transported to SGs (Ansari and Haqqi, 2016; Adjibade et al., 2015). Preliminary results of the composition of SGs in animal cells showed that the majority of mRNAs are targeted to SGs. This process occurs with varying efficiencies ranging from <1 to >95% for individual transcripts (Khong et al. 2017). Currently, the involvement of SGs in the regulation of the translation process has not yet been clearly determined. The formation of SGs appears to be a consequence of translation inhibition rather than its cause. In G3BP1/2 depleted cells, which do not form SGs, it has been shown that translation is still inhibited during stress (Kedersha et al., 2016; Mateju et al., 2020). Nevertheless, there are reports indicating the presence of the 60S subunit in SGs. Localization studies of L37 and L5 proteins have shown colocalization with SGs (Kimball et al., 2003). In addition, 60S accumulates in stress granules leading to disruption of the protein quality control machinery (Seguin et al., 2014). These data require further elucidation in animal and plant cells.

Recently, a new level of gene expression regulation has been demonstrated, indicating that nucleotide modification in mRNA is involved in this process. This phenomenon has been termed epitranscriptomic regulation. The most common modification found in mRNA is N6-methyloadenosine (m6A). Not fully understood is the role of m6A in regulating transcript localization. Anders et al. (2018), showed that in animal cells, under conditions of oxidative stress, the level of m6A increased in individual transcripts and the total amount of methylated transcripts increased. Interestingly, 96% of the transcripts in which m6A was upregulated in response to stress were located in the SGs. Analysis of methylation sites showed that the presence of m6A near the 5’ ends is a specific mechanism responsible for localizing them to the SGs. The removal of this methylation site in a single transcript results in it not being targeted to the SGs. Ries et al (2023) showed in a knockout cell line of the METTL3 gene (responsible for methylation) that long mRNAs accumulate in the SG as a result of m6A methylation. Long exons induce mRNA methylation and provide scaffolds for numerous protein interactions that direct transcripts to the SGs. The results of animal cell experiments do not clearly indicate the involvement of m6A in mRNA localization in SGs. Analysis of mRNA accumulation in stress granules in WT and ΔMETTL3 mES (without m6A modification) cells, did not show differences in the localization of 13 m6A-modified mRNAs in SGs (Khong et al., 2022). The measurement of m6A levels in mRNA purified from SGs showed that m6A levels were approximately 50% higher than in total cytoplasmic mRNA (Ries et al., 2019). Far fewer results have been published on the role of m6A in plants (Scutenaire et al., 2018, Prall et al 2023). It has been shown that methylation of the heat-activated retrotransposon Onsen in *A. thaliana* leads to localization in SGs and thus suppression of its mobility (Fan et al., 2023). The potential role of m6A modifications in mRNA localization in SGs may be evident by the fact that YTH domain-containing proteins, responsible for m6A recognition, were found in *A. thaliana* SGs (Scutenaire et al., 2018, Kosmacz et al., 2019). In animals, knockout of the YTHDF3 protein resulted in loss of m6A co-localization signal with SGs (Anders et al. 2018). Whether these proteins and level of m6A affect transcript sequestration in SGs requires further study in plants.

In this study, we discovered a new type of SGs with different morphology and determined the roles of epitranscriptomic mechanisms in the functioning of SGs in plants subjected to hypoxia stresses.

## Results

### Formation of stress granules in the roots of *Lupinus angustifolius* during hypoxia

Localization of poly(A) RNA in roots meristematic cells of *Lupinus angustifolius* subjected to hypoxia and reoxygenation was performed (Fig. 1). In the cytoplasm the different size poly(A) RNA structures during stress were observed. To verify whether the structures could represent stress granules, we performed a localization of the marker protein of the SGs, i.e. PAB2 (Fig. 1A-S). In all stages of hypoxia, strong colocalization of cytoplasmic aggregates, poly(A) RNA and PAB2 was observed (Fig. 1D-O). Since the formation of SGs is inhibited when translation elongation is blocked, we treated roots during hypoxia with cyclohexamide (Fig. S1). The drug prevent assembly of cytoplasmic aggregates. Therefore the cytoplasmic structures with poly(A) RNA and PAB2, which formed in lupin during hypoxia, constitute SGs. Next, we analyzed SGs formation during the successive stages of hypoxia. After 1 hour, round and similar in size SGs emerged in the cytoplasm (Fig 1D-F). In the next stage their area increased significantly, but the number of SGs decreased (Fig. 1T,U). They were elongated, composed of two or more circular structures (Fig. 1G-I). This strongly suggests the fusion of single SGs into larger structures after 6h of hypoxia. The fusion of SGs continued in the cells of roots exposed to hypoxia stress for 9h and 15h (Fig. 1J-O). Successive hours of hypoxic stress slightly increased the number of SGs (Fig. 1T). After stress removal and culturing seedlings for 6h in optimal oxygen concentration (reoxygenation) the SGs were not observed any longer (Fig. 1P-S). The first hour of stress resulted in a high level of poly(A) RNA in the cytoplasm and the emergence of small stress granules containing 3.3% of transcripts (Fig. 1V). However, in 6h and 9h stressed root cells, the quantity of poly(A) RNA increased in SGs. Experimental 15h hypoxia caused the highest poly(A) RNA accumulation in SGs, a 20.4% of the total cytoplasmic pool (Fig 1V). The stages of SG formation consist of the appearance of small structures, which then merge into larger ones, which in turn leads to a decrease in their number. We also showed that the process of poly(A) RNA accumulation in the SGs increases and is proportional to the duration of the stress and is accompanied by a decrease in the number of transcripts in the cytoplasm (Fig. G-O). This indicates that the poly(A) RNA in SGs originated in the cytoplasm.

**Fig. 1.**
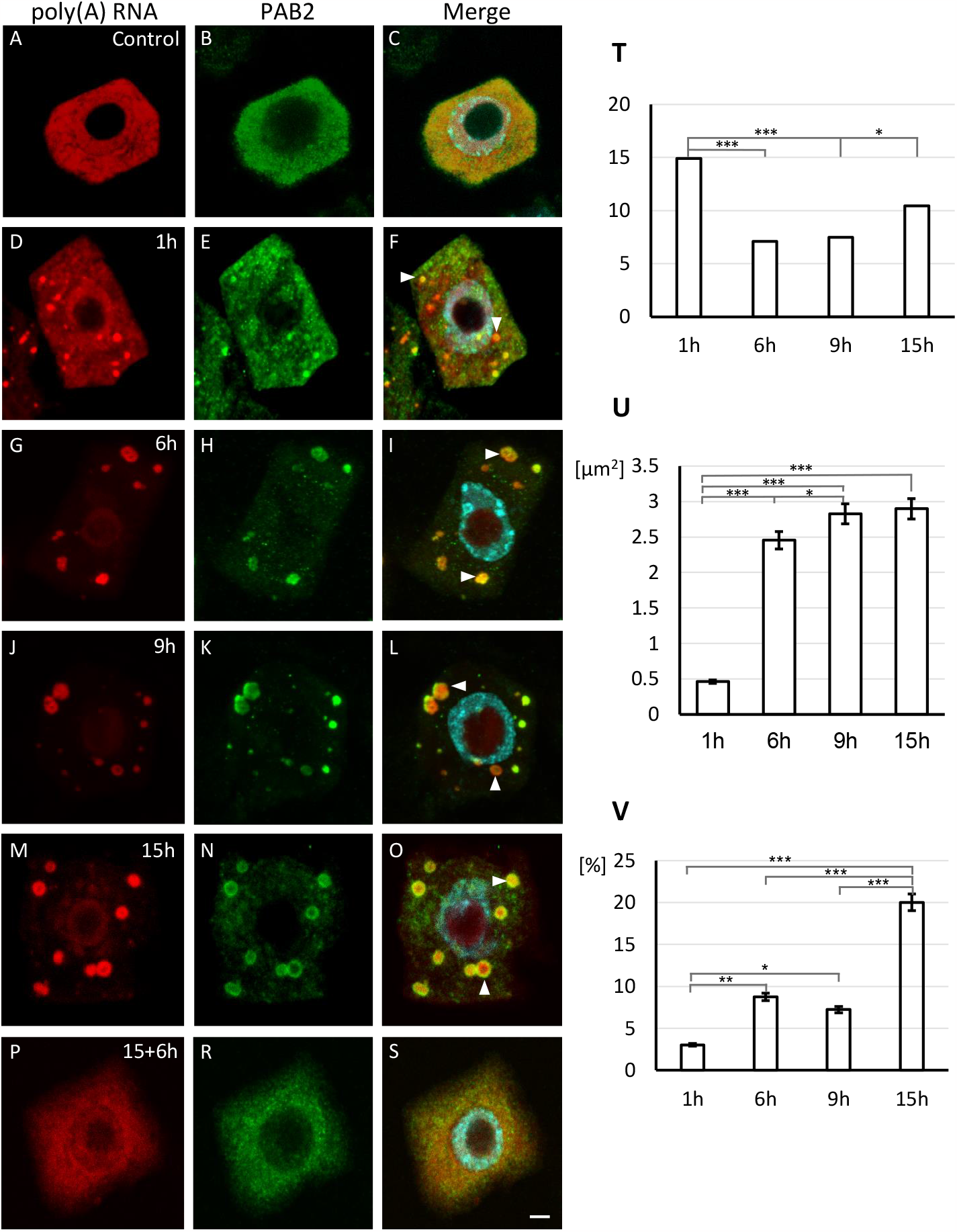
Localization of poly(A) RNA (red fluorescence) and SGs marker protein PAB2 (green fluorescence) in meristematic cells of *L. angustifolius* roots during normoxia (**A-C**), hypoxia (1, 6, 9, 15 h; **D-O**) and reoxygenation (15 h hypoxia followed by 6 h reoxygenation; **P-S**), merge of signals and DAPI staining (**C, F, I, L, O, S**), bar 10 μm. During deprived oxygen conditions, the accumulation of poly(A) RNA was observed in SGs (arrowheads). Disassembly of SGs occurs after the removal of stress. Quantitative analysis of amount of SGs (**T**), size of SGs (**U**) and the percentage of cytoplasmatic pool of poly(A) RNA in SGs (**V**) during hypoxia (1, 6, 9, 15 h) stress. Student’s t test was done to assess significance, with *, P ≤ 0.05; **, P ≤ 0.01; and ***, P ≤ 0.001. Error bars represent standard deviations.

### Bi-Zonal Stress Granules are in roots of Lupinus angustifolius

The PAB2 and poly(A) RNA are not homogeneously distributed in the SGs. To confirm this observation, FISH and immunochemical reactions on resin sections were performed. Both poly(A) RNA and PAB2 were in the periphery of SGs, forming a ring with space in the center free of the tested molecules. These types of SGs were primarily observed after 6-15 hours of hypoxia (Fig. 2 A-F). PAB2 formed fine clusters and poly(A) RNA had a more uniform distribution in the ring. In most of the observed SGs, the central area after staining RNA was smaller than with protein. This may suggest the presence of a small amount of poly(A) RNA in the center of SGs. Subsequently, the SGs were analyzed at electron microscope. The stress granules found in hypoxia lupin roots were two-zone, consisting of a ring formed of coiled dense fibers, and the central brighter area (Fig. 2G). SGs presenting this morphology have not been described in plants yet.

**Fig. 2.**
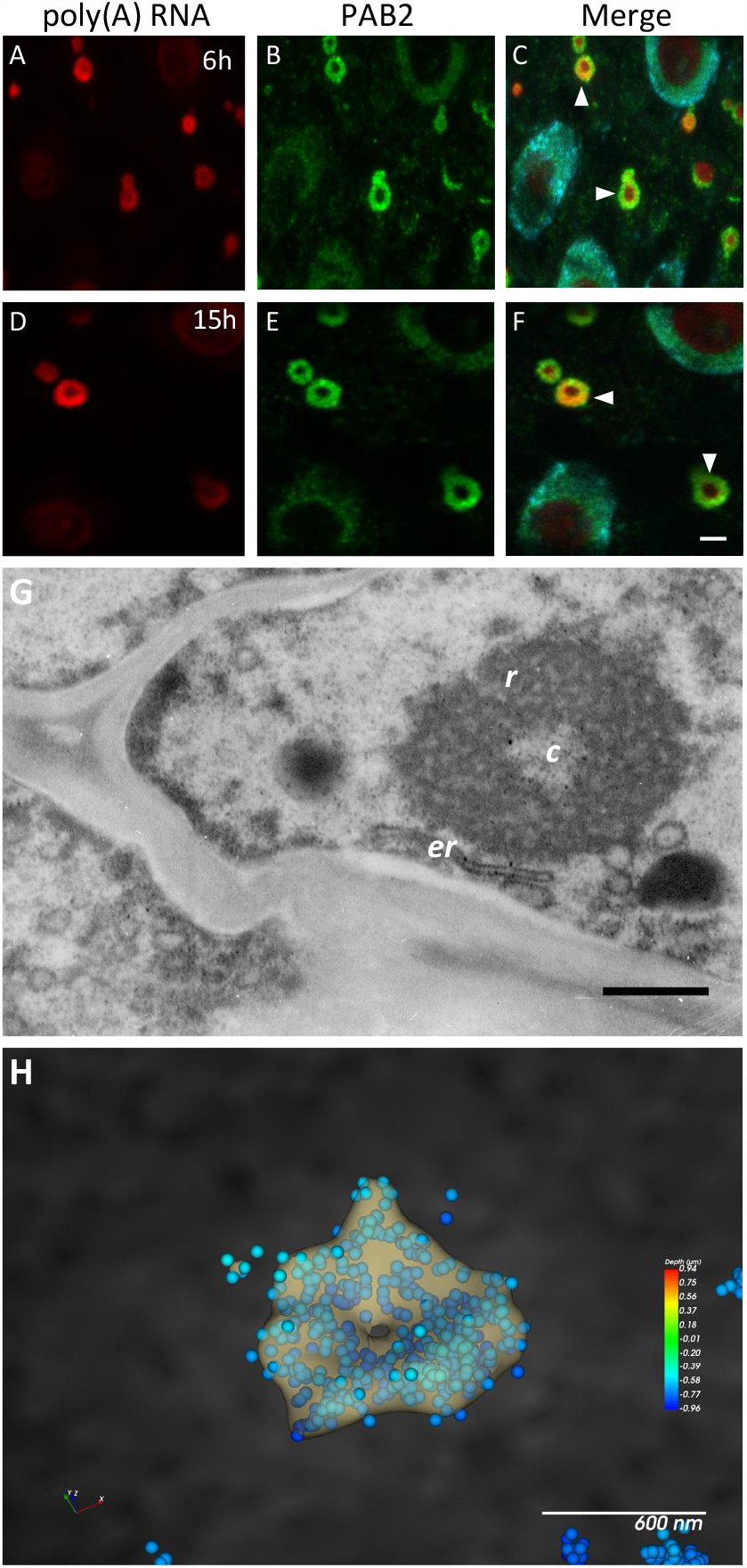
Localization of poly(A) RNA (red fluorescence) and SGs marker protein PAB2 (green fluorescence) in resin sections of meristematic cells of *L. angustifolius* roots in normoxia (**A-C**) and 15 h of hypoxia (**D-F**), merge of signals and DAPI staining (**C, F**), the arrowheads indicate SGs, bar 5 μm. Ultrastructural analysis of bizonal SGs (**G**), *r* coiled dense fibers ring, *c* central brighter area, *er* endoplasmatic reticulum, bar 3 μm. STORM analysis (**H**). The dots represent the poly(A) RNA individual molecules that were detected during the acquisition. Dots are color-coded by depth. Dot size is in the image is set to 50 nm. The SG structure in the image is the cluster’s isosurface calculated by performing a Cluster analysis on the data set, bar 600 nm.

In the next step we used STORM, a high-resolution confocal microscope to analyze distribution of poly(A) RNA. The dots represent the individual localizations of molecules that were detected during the acquisition. The signal was distributed uniformly over the ring of SGs and it didn’t occur in the center of this structure, confirming the two-zone nature of SGs (Fig. 2H). To summarize, SGs in *Lupinus angustifolius* consist of two areas of a different composition and ultrastructure.

SGs were often surrounded by endoplasmic reticulum at EM level (Fig. 2G). The structure of the central zone reassembles cytoplasm outside the SGs. To study the nature of this structure we performed localization of 18S and 26S rRNAs i.e. small and large subunits of ribosomes, respectively (Fig 3A-L). In the center of SGs 26S RNA was observed (but not in the ring-like area). Interestingly, compared to other parts of cytoplasm, higher concentration of 26S was detected around the SGs (Fig. 3D-F). A similar distribution (but without concentration around SGs) was observed for 18S rRNA (Fig. 3 J-L). Quantitative rRNAs analysis revealed that the level of both studied ribosomal transcripts was similar in the cytoplasm and in the center of SGs (Fig. 3M, N). The above results suggest that ribosomes are not localized within the ring of SGs but only in the central region of SGs, and their arrangement and abundance are comparable to or higher than those observed in the cytoplasm.

**Fig. 3.**
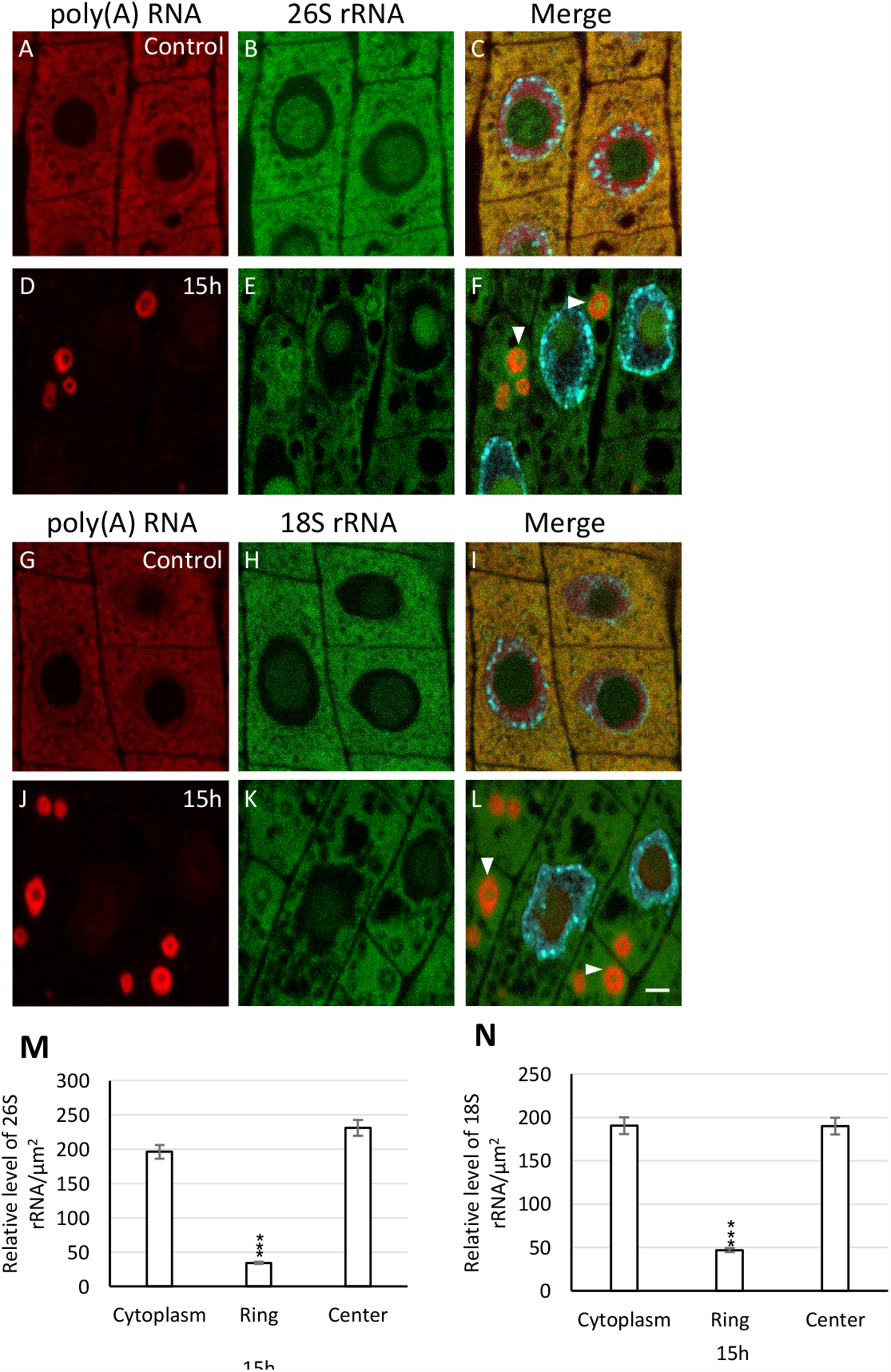
Localization of poly(A) RNA (red fluorescence) and 26S and 18S rRNA (green fluorescence) in meristematic cells of *L. angustifolius* roots in normoxia (**A-C, G-I**) and 15 h of hypoxia (**D-F, J-L**), merge of signals and DAPI staining (**C, F, I, L**), the arrowheads indicate SGs, bar 10 μm. Quantitative analysis of 26S (**M**) and 18S (**N**) rRNA in the cytoplasm and in two zones of SGs: ring and central area. Student’s t test was done to assess significance, with ***, P ≤ 0.001. Error bars represent standard deviations.

### RNA-Seq reveals transcriptome of lupine is deeply reshaped in hypoxia

To find out differentially expressed poly(A) RNAs in hypoxia, we performed high-throughput sequencing of *L. angiustifolius* roots in three conditions: hypoxia, reoxygenation and normoxia. To infer systematic patterns across biological replicates, principal component analysis (PCA) was performed (Fig. S2A). The clustering has shown limited biological variation among the triplicates, with strong distinction between the compared groups, confirming the plant’s remarkably strong response to investigated experimental factors (Fig. S2B). A similar number of genes were shown to be up- and down-regulated (Fig. 4A-C). The largest number of differentially expressed genes (DEGs) is observed between hypoxia and normoxia (21036), confirming the lupine’s strong reaction to the stress of hypoxia. Importantly, the DEGs in that group contain a number of Hypoxia Responsive Genes (HRG), previously identified in *A. thaliana* (Mustroph et al. 2010). All of the four genes marked on the vulcano plot (ADH1, HUP7, PDC1, HRA1) belong to this group get upregulated in hypoxia, with the fold change between 5.8 and 8.9 (Fig. 4A). Interestingly, among the upregulated DEGs there are m6A-metabolism related genes as well, such as the following m6A writers: MTB, Vir, and HAKAI (Fig. 4A). Not all m6A-linked genes behave the same; for instance demethylase (eraser) ALKBH9B gets upregulated, while its close homolog, ALKBH9C, displays lowered expression levels in stress (Fig. 4A). Of note, also housekeeping genes have lower expression during stress, as exemplified by L44, L37, and RPB1. When it comes to reoxygenation compared to hypoxia, the observed patterns are quite different (Fig. 4B). In turn reoxygenation vs. normoxia HRG genes are still active after stress is removed but substantially less than in hypoxia. The erasers are less expressed compared to normoxia (Fig. 4C). Expression levels of some (Rpb1) but not all (L37, L44) housekeeping genes are elevated compared to normoxia.

**Fig. 4.**
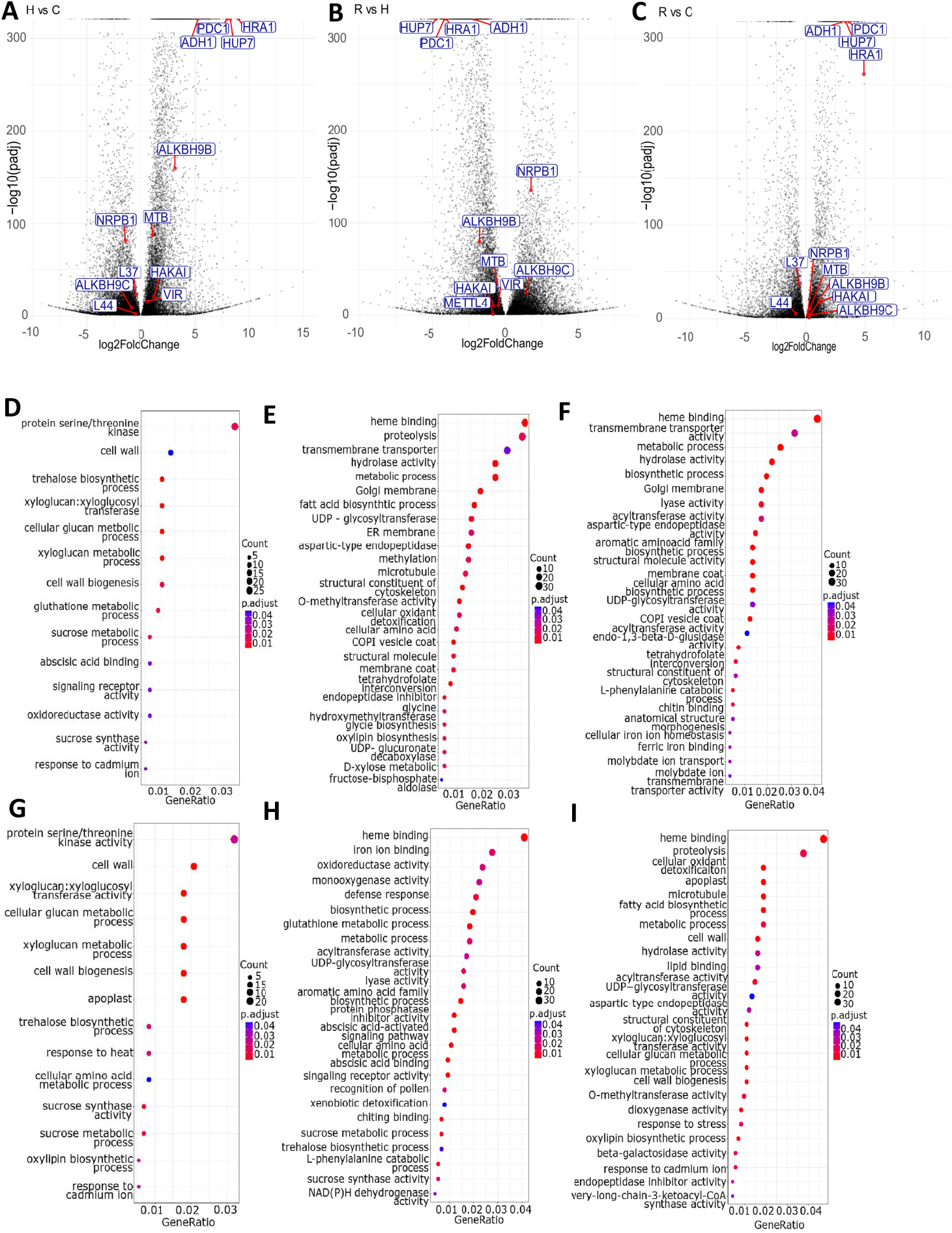
A volcano plot of identified DEGs for hypoxia vs normoxia (**A**), reoxygenation vs hypoxia (**B**) and reoxygenation vs normoxia (**C**) comparisons. Indicated are Hypoxia Responsive Genes (ADH1, HUP7, PDC1, HRA1), m6A-metabolism related genes (m6A writers: MTB, Vir, and HAKAI; m6A erasers: Alkbh9b, Alkbh9c) and housekeeping genes (L37, L44, RPB1). A dotplot representation of Gene Ontology terms overrepresented among up- (**D**,**F**,**H**) and down- (**E, G, I**) regulated genes in hypoxia vs normoxia (**D, E**), reoxygenation vs hypoxia (**F, G**), reoxygenation vs normoxia (**H, I**) comparisons respecitvely.

To comprehensively explore the regulatory networks involved in the hypoxia response, we conducted Gene Ontology (GO) analysis on sets of upregulated and downregulated genes during stress compared to normoxia. The upregulated DEGs were categorized into 24 GO terms (Fig. 4D). The primary six subnetworks include: 1. protein serine/threonine kinase activity; 2. regulation of cell wall including xyloglucan 3. trehalose biosynthetic process, 4. sucrose metabolism, 5. ABA response and 6. oxidoreductase activity. Among the downregulated genes, several categories associations with various cellular functions such as organelle function, cytoskeleton, and vesicle transport, specifically in: 1. heme binding, 2. proteolysis, 3. transmembrane transporter activity, 4. cellular oxidant detoxification (Fig. 4E).

Subsequently we conducted a distinct Gene Ontology (GO) analysis for both upregulated and downregulated genes in reoxygenation vs. hypoxia (Fig. 4F,G). The upregulated genes are in 25 categories, including genes involved in heme binding, transmembrane transporter activity, metabolic process, structural constituent of cytoskeleton (Fig. 4F). In turn, genes with lowered transcriptional activity are also involved in the regulation of heme binding and proteolysis, and in the fatty acid biosynthetic process, response to stress (Fig. 4G). The GO analysis of reoxygenation vs. hypoxia revealed several contrasting trends compared to hypoxia vs. normoxia, indicating a robust recovery after the stress period. Notably, all comparisons underscored strong regulation of biogenesis and cell wall components (Fig. 4D-I).

Next we selected several genes shown to be up- or downregulated by the RNA-Seq data for validation by qRT-PCR. Of the HRGs known from the *A. thaliana* ADH1, HUP7, PCO1 and WIN1 were examined (Fig. 5A,B). For the first two, a high increase of transcripts during hypoxia was observed, which was consistent with RNA-seq results. In turn, reoxygenation resulted in a reduction in the amount of mRNAs (Fig. 5A,B). During stress the expression of PCO1 did not change and WIN1 strongly decreased, indicating that these homologues did not responded to oxygen deprived conditions in lupine (Fig. 5B). In contrast, the transcriptional activity of L37, L44 and RPB1 decreased during hypoxia and increased after stress removal, confirming that genes were not responsive to oxygen reduction in lupine (Fig. 5B). The qPCR results strongly confirmed the RNA-seq data.

**Fig. 5.**
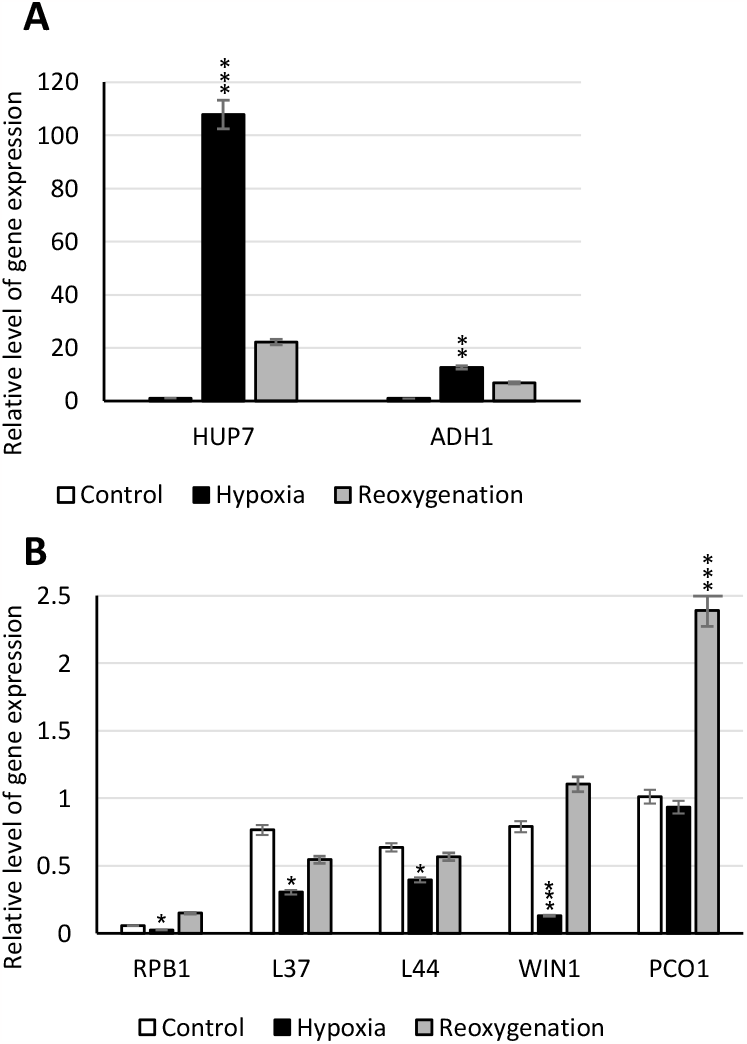
Quantification of expression levels of hypoxia marker genes (**A**) and housekeeping genes (**B**) in normoxia, hypoxia and reoxygenation. Quantitative data were normalized with the expression of UBC5. Data are means of 3 independent biological replicates. Bars represent standard errors. Asterisks indicate statistically significant differences ; *P < 0.05; ***P < 0.001 by Student’s t-test.

### Localization m6A and mRNA in stress granules

Firstly, we conducted fluorescent in *situ* hybridization (FISH) to transcripts, with an emphasis on SGs. The quantity of ADH1 mRNA increased in cells within the first few hours of stress (Fig. 6D-O), but the spatial correlation of the transcripts with SGs could only be estimated after 9 hours of hypoxia (Fig. 6J-L). Within stress granules, three different probes to ADH1 mRNA demonstrated a non-uniform distribution, with its presence predominantly observed only in the central region of structure while being absent from the ring-like. Similar distribution in SGs was observed after 15h of stress (Fig. 6M-O). Quantitative measurements of ADH1 mRNA in the SGs zones and cytoplasm revealed that the ring-like region contained more than four times less mRNA compared to the central region and cytoplasm. The insignificant amount of ADH1 mRNA in the ring-like zone is evidenced by the fact that the level was similar to the amount in the cytoplasm under control conditions. In turn, the amount of transcript in the central part of SGs was only approximately 15% lower than in the cytoplasm cells exposed to 15h stress (Fig. 6T). The distribution of ADH1 mRNA in the SGs and cytoplasm was similar to rRNA. During reoxygenation, SGs disappear, and the amount of ADH1 transcript in the cytoplasm decreased (Fig. 6P-S). Subsequently localization of transcripts of housekeeping genes was performed. The qPCR and RNA-seq data for RPB1 were confirmed by FISH with probes to the middle sequence of transcript (Fig. 7A-S). Those methods have shown that the level of transcripts lowers during hypoxia and strongly increases after stress removal. SGs with RPB1 mRNA were observed at 6 hours of hypoxia (Fig. 7G-I), and the ratio of mRNA content between SGs and the cytoplasm consistently increased up to 15 hours of stress (Fig. 7W). In contrast to mRNA ADH1, the distribution of RPB1 within SGs appeared to be relatively uniform. The level of RPB1 mRNA in the SGs was almost twice as high as in the cytoplasm during long-term hypoxic stress (15h) (Fig. 7W). This suggests that the decrease in transcript levels in the cytoplasm correlates with their accumulation in SGs. Upon reoxygenation dispersal of these structures was accompanied by a strong signal in the cytoplasm (Fig. 7P-S). A similar localization pattern was observed for other transcripts, HUP7 hypoxia response gene and 4 housekeeping genes (L37, L44, PCO1, WIN1) (Fig S3, 4). Additionally, the localization of the 5’ UTR of RPB1 mRNA was conducted (Fig. 7T-V). Interestingly, a slightly different distribution was obtained using a probe recognizing the middle RPB1 mRNA sequence. The 5’ UTR of RPB1 was exclusively found in the ring of SGs, as the probe to poly(A) RNA. This suggests distinct positioning of the 5’-ends, 3’-ends, and the middle sequences of the transcripts. The mRNA ends were in the ring of SGs, where there was no rRNA, and the middle of transcripts were in the central area of SGs. Recent studies have demonstrated the presence of m6A-binding protein ECT2 in SGs of *A. thaliana* (Kosmacz et al., 2019, Fan et al., 2023). Hence, we aimed to investigate whether RNAs with m6A is present in SGs of lupine. Dynamic changes in the level of m6A in SGs were observed in subsequent hours of hypoxia (Fig 8A-O). During the first hours (3-6h) of hypoxia m6A accumulated or entirely co-localized with SGs (Fig 8D-I). At successive stages of hypoxia, a reduction in m6A within stress granules was noted (Fig. J-O). The ratio of cytoplasm to SGs for m6A was highest at the onset of stress and decreased by half during the 9-15 hours of hypoxia (Fig. 8P). This suggests the accumulation of m6A-rich RNA in the early stages of stress granule assembly and their reduction during stress. To investigate the relationship between the level of m6A and the localization and expression levels of transcripts, we employed the Met-RIP (Methylated RNA Immunoprecipitation) technique. The results obtained indicate substantial variations in m6A levels among mRNAs (Fig. 9). We observed an increase in m6A during hypoxia and decrease during reoxygenation for all transcripts. However, both ADH1 and HUP7 mRNAs exhibit significantly lower m6A quantities at all stages compared to the other genes (Fig. 9). The highest expression levels of these genes were observed during hypoxia. It is noteworthy that a higher increase in m6A levels during hypoxia, compared to normoxia (Fig. 9), correlates significantly with the decreased amount of mRNA among housekeeping genes (Fig. 5B). This relationship was most visible for WIN1 and L37. Obtained results indicate a clear negative correlation between the m6A level and the expression of the studied transcripts. This correlation is observed across the examined genes and analyzed stages. An additional transcript with a high level of m6A, accumulated in stress granules (SGs). In turn, HUP7, despite its equally low level of m6A, was present in both SGs and the cytoplasm. Notably, the expression of HUP7 was the highest among all analyzed mRNAs during hypoxia, surpassing even ADH1.

**Fig. 6.**
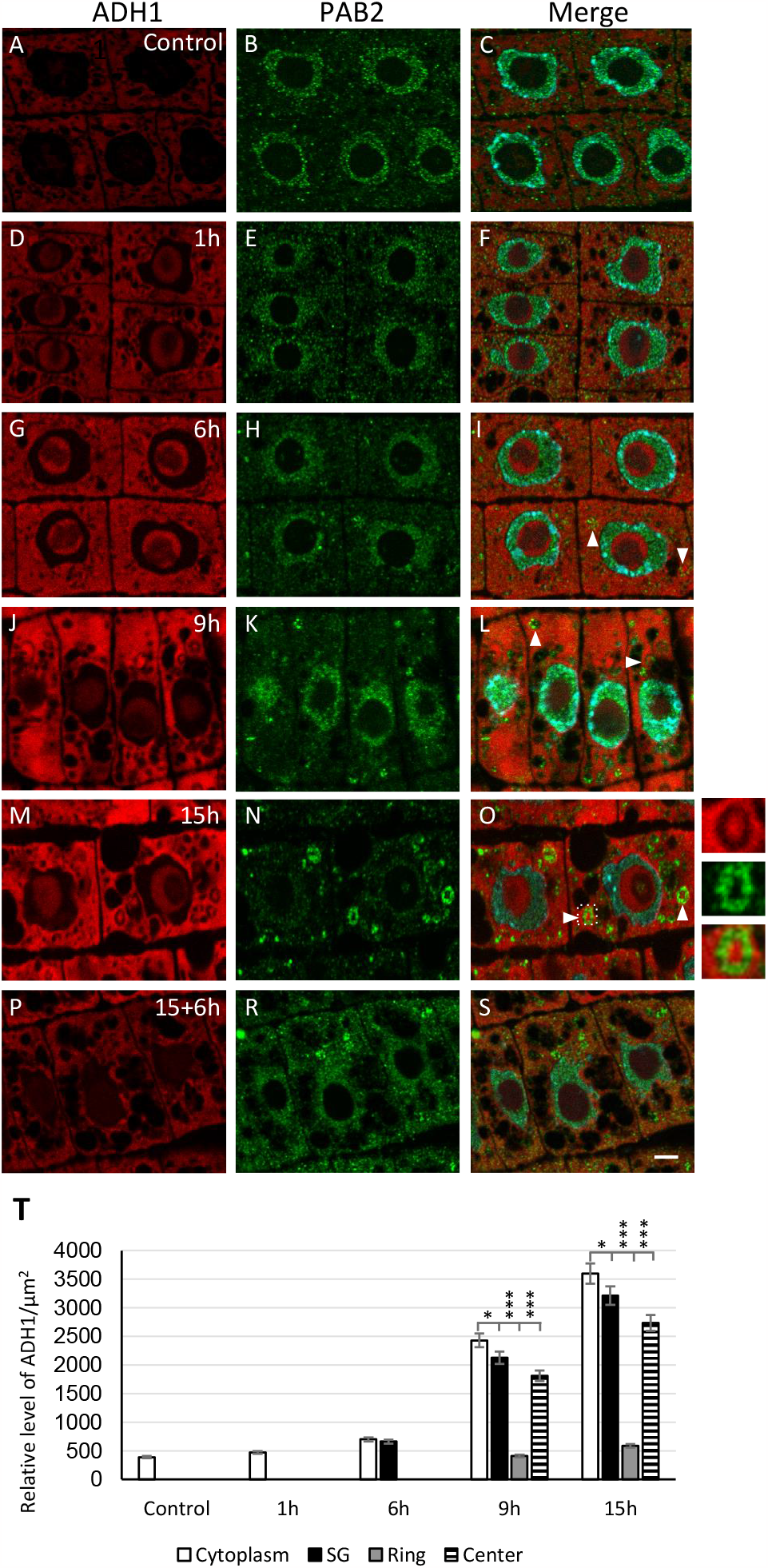
Localization of ADH1 mRNA (red fluorescence) and SGs marker protein PAB2 (green fluorescence) in meristematic cells of *L. angustifolius* roots in normoxia (**A-C**), hypoxia (1, 6, 9, 15 h; **D-O**) and reoxygenation (15h hypoxia followed by 6 h reoxygenation; **P-S**), ADH1 transcripts in central zone of SGs are visible (arrowheads), the right-hand panel represents the magnification of SG which is marked with a square, merge of signals and DAPI staining (**C, F, I, L, O, S**), bar 10 μm. The relative fluorescence intensity of ADH1 mRNA in the cytoplasm, SGs and two zones of SGs: ring and central area in control conditions and during hypoxia (T). Student’s t test was done to assess significance, with *, P ≤ 0.05; ***, P ≤ 0.001. Error bars represent standard deviations.

**Fig. 7.**
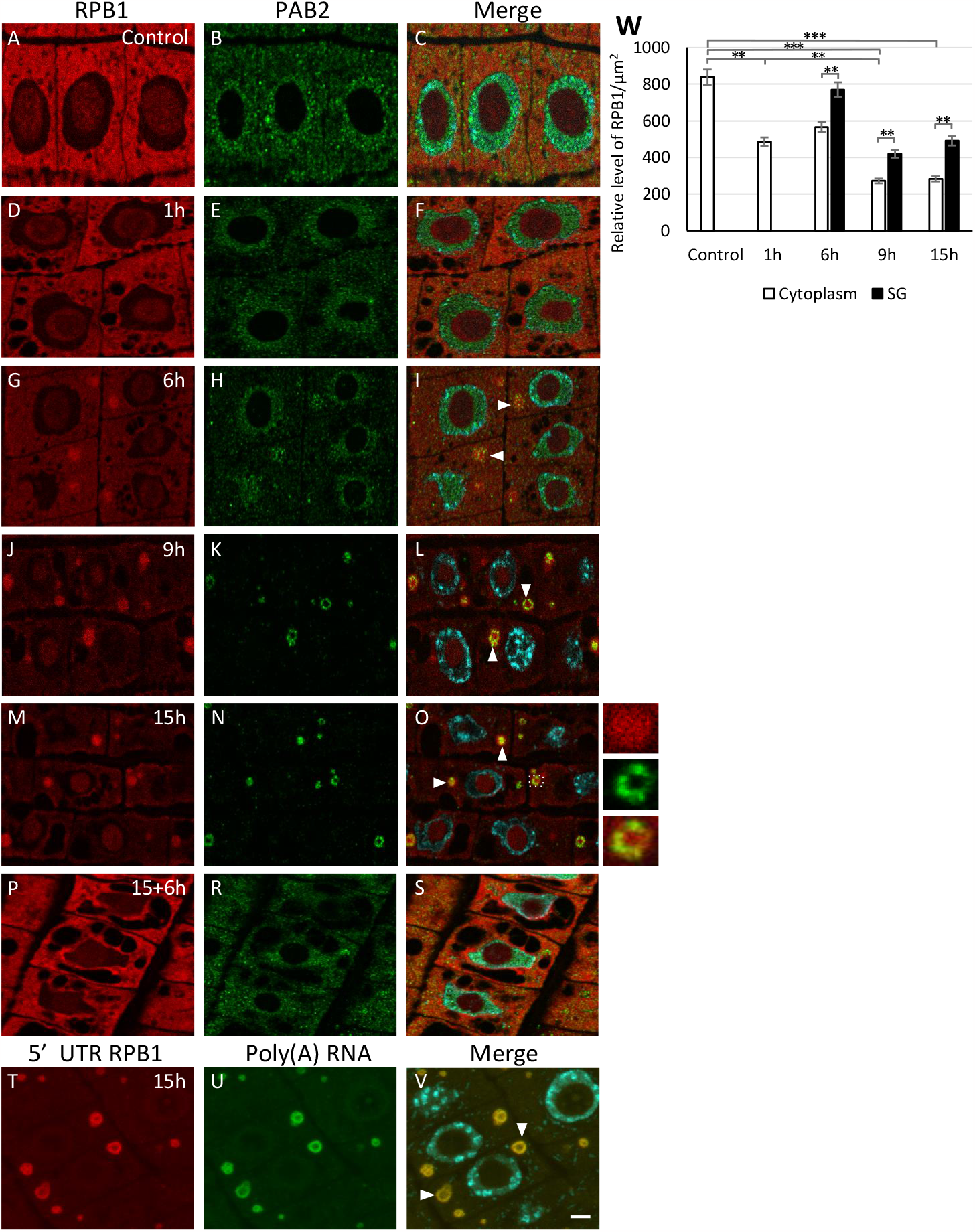
Localization of RPB1 mRNA (red fluorescence) and SGs marker protein PAB2 (green fluorescence) in meristematic cells of *L. angustifolius* roots in normoxia (**A-C**), hypoxia (1, 6, 9, 15 h; **D-O, T-V**) and reoxygenation (15 h hypoxia followed by 6 h reoxygenation; **P-S**), homogenous distribution of RPB1 transcripts detected by FISH using probe to the middle sequence of mRNA are observed in all area of SGs (arrowheads, **I, L, O**) while using probe to 5’ UTR RPB1 transcripts in ring of SGs are visible (arrowheads, **V**), the right-hand panel represents the magnification of SG which is marked with a square, merge of signals and DAPI staining (**C, F, I, L, O, S**), bars 10 μm. The relative fluorescence intensity of RPB1 mRNA in the cytoplasm and SGs during normoxia and hypoxia (**W**).

**Fig. 8.**
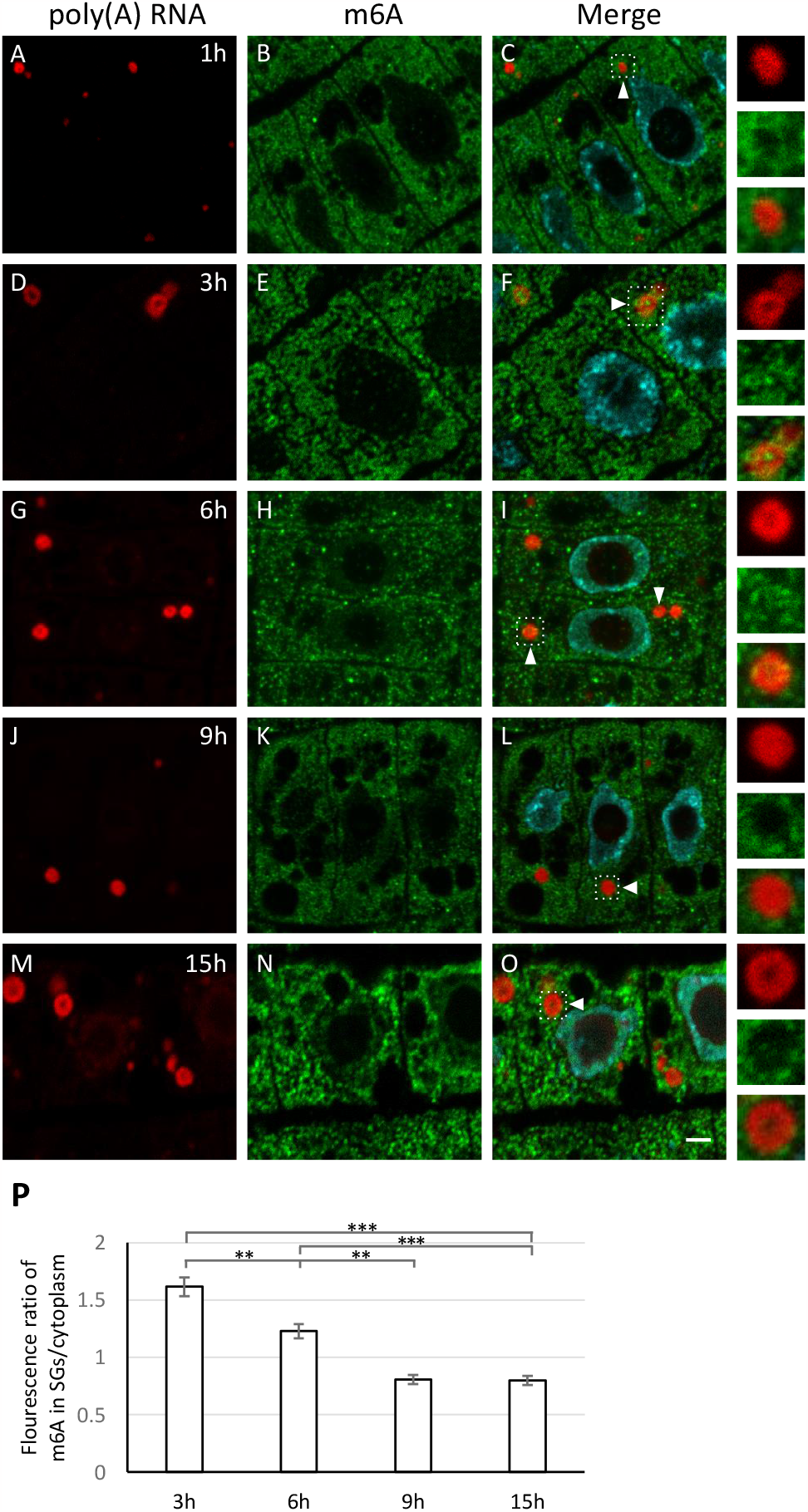
Localization of poly(A) RNA (red fluorescence) and m6A (green fluorescence) in meristematic cells of *L. angustifolius* roots in hypoxia (**A-O**), during 3-6 h of hypoxia (**D-I**) accumulation of m6A in SGs is observed while at next successive hours of hypoxia (9-15 h, **J-O**) a reduction of m6A in SGs is visible (arrowheads), the right-hand panel represents the magnification of SG which is marked with a square, merge of signals and DAPI staining (**C, F, I, L, O**), bar 10 μm. Analysis of SGs/cytoplasm ratio for m6A during 3, 6, 9, 15 h hypoxia conditions (**P**).

**Fig. 9.**
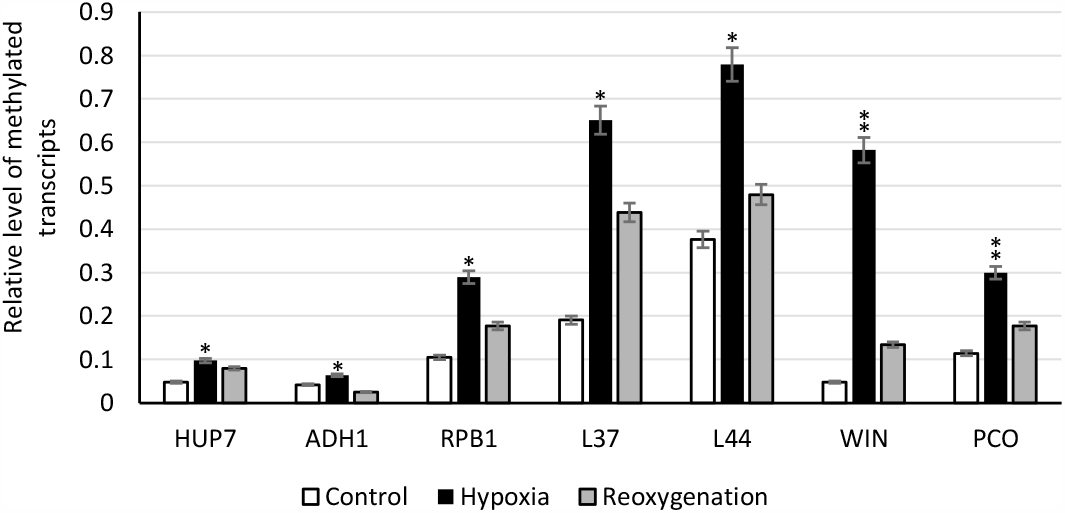
Quantitative measurements of methylation (m6A modification) in HUP1, ADH1, RPB1, L37, L44, WIN1, PCO1 transcripts in *L. angustifolius* roots subjected to control (normoxia), hypoxia and reoxygenation.

Mutants of *L. angustifilius* are not available, therefore, in order to study the function of m6A in the formation of SGs, we used the mtaABI3:MTA *A. thaliana* mutant. The ABI3 promoter drives the strong embryo expression of MTA resulting in a high level of m6A, enabling plant embryo lethality to be bypassed. However, this promoter drives a very low level of expression post-germination (in 14 – days old seedlings), giving rise to plants with 80 to 90% less amounts of m6A in RNA than their WT counterparts. We crossed lines mtaABI3:MTA with ECT2-GFP (reader of m6A occurred in SGs) and Rpb47b-GFP (marker of SGs). Microscopic analysis has shown lack of accumulation of ECT2 in cytoplasm of plants with decreased levels of m6A (Fig. 10A-F). Interestingly, poly(A) RNA clusters resembling SGs were observed in the cytoplasm. It’s suggested that ECT2 is not a crucial protein for assembly. SGs occurred in mtaABI3:MTA x Rbp47b (Fig. 10 G-J), but the number was approximately 25% lower than in WT (Fig. 10L). The possibility of m6A participating in the formation of SGs was evidenced by a slightly lower level of poly(A) RNA in mutants with lower levels of m6A (Fig. 10K). However, as in *L. angustifolius*, in SGs of *A. thaliana* the m6A level in long-term hypoxia was relatively low compared to cytoplasm (Fig. S5). This suggests an involvement of m6A in the first stages of formation of SGs.

**Fig. 10.**
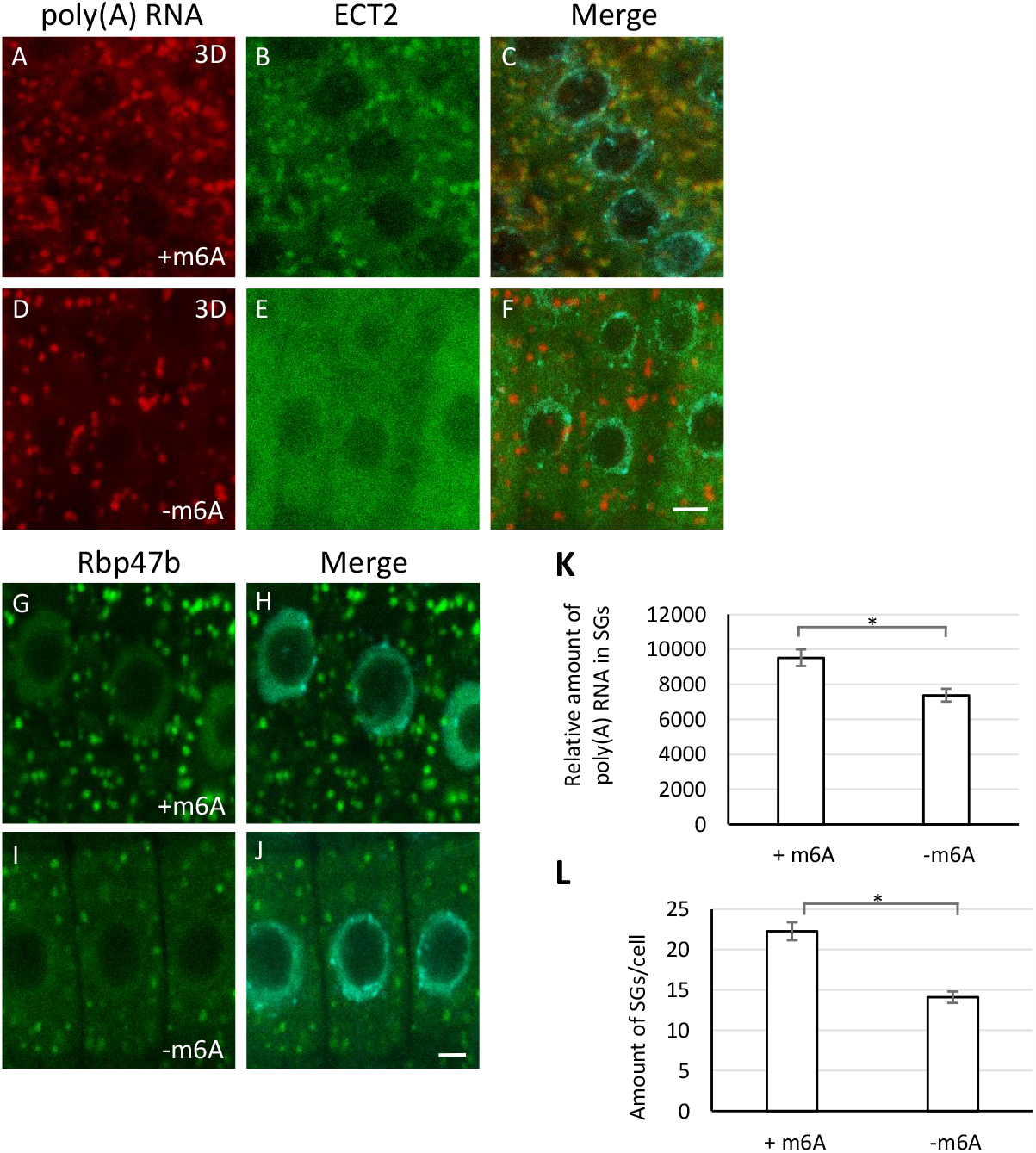
Detection of poly(A) RNA (red fluorescence) and ECT2-GFP protein (green fluorescence) in *A. thaliana* roots cells of ECT2-GFP (**A-C**) and mtaABI3:MTA x ECT2-GFP (**D-F**) mutants and Rpb47b-GFP protein (green fluorescence) in Rpb47b-GFP (**G-H**) and mtaABI3:MTA x Rpb47b-GFP (**I-J**) mutants during long-term hypoxia, merge of signals and DAPI staining (**C, F, H, J**), bar 10 μm. Measurements of the relative level of poly(A) RNA in SGs (**K**) (ECT2-GFP and mtaABI3:MTA x ECT2-GFP) and the amount of SGs (**L)** (Rbp47b-GFP and mtaABI3:MTA x Rpb47b-GFP) in roots cells of mutants subjected to hypoxia conditions. Student’s t test was done to assess significance, with *, P ≤ 0.05; Error bars represent standard deviations.

## Discussion

We precisely documented emergent of the SGs in crop plant e.i. *Lupinus angustifolius* during hypoxia stress. These structures are formed by nucleation of poly(A) RNA and PAB2 and then fusion into larger SGs. This is consistent with data on the number and size of SGs in *A. thaliana*. Long heat-stress caused a decrease in the number and a slight increase in the volume of the analyzed structures (Gutierrez-Beltran et al., 2015; Hamada et al., 2018). What definitely differs between these structures in both models is the diameter and shape. In lupine, SGs are few times larger than in *A. thaliana* and consist of two zones: a ring and a central area. These areas differ in the composition and function. The first one is larger and consists of coiled fibers rich in poly(A) RNA, PAB2 and all the studied transcripts, except ADH1. The distribution of molecules in the ring-like part is homogeneous. Analysis of poly(A) RNA under a high-resolution microscope did not reveal any clusters that could resemble cores of SGs. In turn, the central part of the SGs lacks the poly(A) RNA tail and the PAB2 protein. Additionally, the analysis of various mRNA fragments revealed that in the ring of SGs there are 5’ RPB1 mRNA and 3’ poly(A) mRNAs, while the middle sequences L37, L44 and RPB1 are present in both parts of SGs. This indicates that anchoring the 5’ and 3’ of mRNA in the SGs ring may inhibit association of small and large subunit of the ribosome to transcripts. The presence only of middle mRNA fragments in the central part where 18S and 26S rRNA are present prevents protein synthesis. This suggestion is confirmed by the presence of ADH1 mRNA only in the central part, the translation of which is necessary to survive hypoxia stress. In animal mRNAs localized in SGs can undergo translation (Mateju et al., 2020) while our research shows that in lupine, where SGs are much larger, regulation of synthesis proteins may be related by the different distribution of 5’-, 3’ – ends and middle sequence of the transcript within SGs.

We studied the possibility of m6A participation in SGs assembly in lupinus under hypoxia. In animal cells N6-methyladenosine (m6A) has been proposed to target transcripts to SGs (Fu and Zhuang, 2020; Ries et al., 2023). These modifications can impact an mRNA’s propensity to form LLPS aggregates or to interact with SG-associated proteins. It has been shown in oxidative stress that levels of m6A modification significantly increase in human cell lines, mainly in the 5’ UTR and 5’ region of the coding sequences of SG-associated transcripts (Anders et al., 2018). In *A. thaliana* model Kosmacz et al. (2019) have shown m6A reader, ECT2, 4 in SGs under heat stress. In turn, ECT2 (YTH9), a well-characterized m6A reader in plants, is re-localized from the cytoplasm to the SGs under heat shock (Scutenaire et al., 2018). Our data indicates that m6A RNAs are present in SGs because we didn’t observe the ECT2-reader of m6A in SGs in mutant of *A. thaliana* with reduction of m6A. In lupinus the mRNAs of housekeeping genes with high level of m6A (MetRIP, whole cells) mostly are present in the SGs during hypoxia stress. This suggests the involvement of m6A in the accumulation of transcripts in SGs. The exception is HUP7, where, despite the low level of m6A, mRNAs are in SGs, but the expression level of this gene is very high and cells show a high amount of mRNA. It has been shown that regardless of its function, RNAs can be accumulated in the SGs (Tong et al. 2022). It may depend on the amount of it as in ER-stress in *A. thaliana*. The decline in translation efficiency results in transcripts of the Unfolded Protein Response (UPR) not being loaded onto polyribosomes in response to ER stress. Instead, the mRNA accumulates in SGs (Kanodia et al., 2020). The ADH1 mRNA with low level of m6A and high but lower than HUP7 expression are out of the SGs. However, due to its high expression level, HUP7 mRNA depleted m6A, is present both in the SG and cytoplasm. The balance between the level of m6A and transcript expression is responsible for the localization of mRNAs in SGs. The measurements of m6A-rich RNA have shown different quantity in SGs. The amount of m6A is relatively high at the beginning of stress, but in lupin quantity of modified nucleotide decreases as the stress conditions are prolonged. In *A. thaliana* we also didn’t observe strong accumulation of m6A in long-term hypoxia. However, in mutants we have shown that lowering the level of m6A in mtaABI3:MTA blocks ECT2 accumulation and reduces the number of SGs in *A. thaliana*. This is also accompanied by a slight reduction in the amount of poly(A) RNA in SGs. This indicates the involvement of m6A in the assembly and composition of the SGs transcriptome in plants, especially in first steps. It seems that being like animals will not be a key factor in the biogenesis of these structures (Khong et al., 2022). Our study suggests that the correlation of the expression of individual transcripts and the deposition of m6A create favorable conditions for formation of SGs in plants.

The observed decrease in the amount of m6A in SGs during stress requires further research, although recent results from Fan et al. (2023) reveal a possible mechanism. It has been shown that the Onsen RNA transposon with m6A is transported to SGs containing the ALKBH9B demethylase in biotic stress. In ALKBH9B mutant an increase in transposon levels was observed in seedlings, including SGs. This work suggests a mechanism to suppress transposon mobility that involves m6A RNA methylation and sequestration of transposon RNA in SGs. The presence of active ALKBH9B in SGs would explain the decrease in the amount of m6A in SGs we observed in subsequent stages of stress. However, Fan et al. (2023) do not reveal a decrease in the amount of m6A in SGs as the biotic stress progresses or the activity of demethylases in these structures. Our results showed that high levels of m6A correlate with lowering quantity in particular/individual mRNAs in the cell. To protect the mRNAs from degradation and store them in SGs until the end of stress, ALKBH9B could demethylated the transcripts. The RNA-seq data support this mechanism in lupin because the level of mRNA ALKBH9B significantly increase in hypoxia and decrease in reoxygenation.

In conclusion, our study identifies two distinct zones within plant stress granules (SGs) with differing functions. Moreover, we shown the limited impact of m6A modification on SGs assembly. Nonetheless, the interplay between m6A modification and the overall transcript abundance in the cytoplasm plays a regulatory role in mRNA partitioning into SGs.

## Materials and methods

### Plant Growth and Treatment

Seeds of *Lupinus angustifolius* cv. Sonet were soaked in 70% (vol/vol) ethanol for 5 min and rinsed five times with sterile, deionized H_2_O. Seeds were soaked in water for 30 min and then germinated for 3 d at 21°C on water-soaked tissue paper in a growth chamber in the dark. After three days of germination, hypoxia was induced in seedlings with 3-4 cm roots by immersing the plants in containers filled with tap water (oxygen concentration in water: 8.7 mg/L). The water level was about 4 cm above the plants and the plants were cultured under the same conditions. Reoxygenation after hypoxia was performed under seed germination conditions.

Translation in *Lupinus angustifolius* roots was inhibited by 4 mg/ml cycloheximide (Merck). After 14h of hypoxia, the seedlings were immersed in water with the inhibitor and transferred to a vacuum chamber for 1h. After this time, some of the roots were fixed, and the remaining roots were transferred to growth chamber with paper soaked in the inhibitor. The roots were treated with inhibitors in the chamber for 6h, 10h, 20h before fixation.

*Arabidopsis thaliana* ecotype Columbia-0, ECT2:GFP, Rpb47:GFP and mtaABI3prom:MTA (Bodi et al., 2012) (kindly provided by Prof. Rupert Fray) were used in this study. Double mutants ABI3:MTA x Rpb47b-GFP and ABI3:MTA x ECT2-GFP were obtained by crossbreeding. After sterilization the seeds were sown on plates containing 2.15 g/L Murashige and Skoog (MS) (Duchefa) medium, 1 ml of Murashige & Skoog x1000 concentrated vitamin mixture: 103.1 mg/l (Duchefa), 10 g/L sucrose, 7.5 g/L agar, pH 5.8 and were incubated at 4 °C for 2 days in dark. The plants were grown in a growth chamber at 21°C under long-day conditions (16 h light/8 h dark cycle). For hypoxia treatment, the MS plates containing the seedlings were submerged for 3 days in a container filled with sterile water (the oxygen concentration in the water: 8.7 mg/L) so that the water level was approximately 5 cm above plants, and the plants were grown in the same conditions but under low light (30–60 μM m2/s).

### In situ specimen preparation

To prepare *Lupinus angustifolius* protoplasts, root meristems were excised and fixed in 4% formaldehyde (Polyscience) in PBS pH 7.2 for 24 h at 4°C. After wash the fixed root tips were placed in citric acid buffered digestion solution pH 4.8 containing 5% cellulase (Onuzuka R-10) and 35 U/ml, pectinase (Sigma) for 2 h at 35°C. After rinsing with PBS pH 7.2, the root tips were squeezed on slides.

To prepare semi-thin sections of *L. angustiofolius* and *A. thaliana* fixed roots were dehydrated in increasing concentrations of ethanol containing 10 mM dithiothreitol (DTT) (Thermo Fisher Scientific), and then embedded in BMM resin (butyl methacrylate, methyl methacrylate, 0.5% benzoyl ethyl ether with 10 mM DTT (Merck)) at -20°C under UV light for polymerization. The material was cut on a Leica UCT ultramicrotome into sections (1.5 μm).

### In situ hybridization and double labeling with m6A or Pab2

After a 1-hour pre-hybridization, hybridization was performed for at least 12 hours at 26°C in hybridization buffer (50% (v/v) (Merck) with 30% (v/v) formamide (Merck) and 1:250 probes. For FISH double labeling of poly(A) RNA and mRNA (coding ADH1, HUP7, PCO1, L37, L44, RPB1), two probes were applied simultaneously. Slides with sections were washed in 2× SSC, and nuclei were stained with Hoechst 33342 (Thermo Fisher Scientific).

For simultaneous localization of RNA and protein, incubations were performed with primary rabbit antibody against N6-methyladenosine (NEB) or PAB2 (courtesy of Cecile Bousquet-Antonelli and Rémy Merret) diluted 1:200 and 1:100, respectively, in PBS pH 7.2 with 1% BSA overnight at 4°C. Slides were washed in PBS pH 7.2 and incubated for 1 hour at 35°C with secondary antibodies diluted 1:250 in PBS buffer with 1% BSA (Alexa Fluor 488-labeled mouse anti-rabbit IgG (Molecular Probes)).

After washing twice in 2x SSC the in situ hybridization was performed as described above. For control reactions probes and primary antibodies were omitted.

### Microscopic analysis and quantitative measurement of fluorescence signal

To calculate the fluorescence signal obtained from the poly(A) RNA in situ hybridization and the diameter of the stress granules, at least 60 cells from three different experiments were analyzed. Between 30 and 60 cells were analyzed for each transcript. For quantitative measurements, each experiment was performed using constant temperatures, incubation times, and concentrations of probes and antibodies. In addition, images were acquired under constant acquisition conditions (laser power, emission band, gain and resolution). The results were recorded using an Olympus FV3000 microscope and FV3000 software, equipped with a DIC H immersion objective with 63x magnification (numerical aperture 1.4) and exposure (400 Hz). Images were acquired sequentially in red (Cy3), green (Alexa Fluor 488) and blue (Hoechst 33342) channels. Optical sections were collected at 0.5 um intervals. Analysis and processing of the obtained images was carried out using ImageJ software. The signal intensity per cell/nucleus/stress granules was measured and expressed in relative fluorescence units or signal/um^2^. Statistical analysis of the quantity antigens was performed using the PAST program To compare all groups and to determine if there were any significant differences between them, Student’s t-test was used.

### Electron Microscopy

For Standard Transmission Electron Microscopy (TEM) a 4-5 mm long roots were fixed in 3% glutaraldehyde (Polysciences) in PBS pH 7.2 overnight at 4°C. After washing in PBS pH 7.2 roots were postfixed in 1% OsO4 in PBS buffer 1h at 4°C. The material was dehydrated in increasing concentrations of ethanol and embedded in Spurr resin (Sigma). Ultrathin section were placed on copper grids, contrasted in 2.5 % lead citrate and 2.5 % uranyl acetate (20 min. each) and examined in TEM (Jeol 1010).

### Single Molecule Localization Microscopy

Bruker Vutara VXL microscope with widefield illumination and Biplane detection for 3D acquisitions was used to analyze the distribution of poly(A) RNA on semi-thin sections of *L. angustifolius* roots. The following parameters were used: 20 ms exposure time, 90% 638 nm laser power, 692/85 nm emission filter for Cy5 channel, 50 background threshold used in Localization algorithm. Cluster Analysis was performed with the following parameters: DBSCAN cluster algorithm, 0,3 μm maximum particle distance, 50 minimum particle count to form a cluster, 50 nm isosurface particle size

### RNA isolation and m6A-RNA immunoprecipitation

Total RNA was isolated from 100 mg L. angustifolius roots using TRIzol reagent (Thermo Fisher Scientific) and Direct-zol RNA MiniPrep Kit (Zymo Research).

For immunoprecipitation of m6A RNA, total RNA was extracted from *L. angustifolius* roots as described above, followed by polyA enrichment using the PolyATtract® mRNA Isolation Systems (Promega). 9 μg of polyA RNA (1 μg/μl) was used for immunoprecipitation. Immunoprecipitation was performed with the EpiMark N6-Methyladenosine Enrichment Kit (NEB) according to the manufacturer’s instructions. Briefly, 25 μl of magnetic Dynabeads protein G for Immunoprecipitation (Thermo Fisher Scientific) were washed with 250 μl of reaction buffer. Next, 2 μl of anti-m6A antibody/per sample was incubated with the beads for 1 hour at 4°C/4rpm. The beads were washed twice with reaction buffer and resuspended in 250 ul of reaction buffer. 9 μg poly(A) RNA was spiked in with 1 μl of positive and negative control from the kit (1:1000 dilution) and made up to 13 μl final volume with Nuclease-free water. As input, 10% of the sample volume (1.3 ul) was taken and the residue was added to the beads. 3 ul of RNasin Ribonuclease Inhibitor (Promega) was added to the beads and incubated for 2 hours at 4°C/3rpm. After incubation, the beads were washed twice with reaction buffer, LSB buffer (0.02M Tris HCl pH 8, 0.002M EDTA, 1% Triton X-100, 0.15M NaCl, 0.1% SDS) and HSB buffer (0.02M Tris HCl pH 8, 0.002M EDTA, 1% Triton X-100, 0.5M NaCl, 0.1% SDS). RNA was extracted using Acid Phenol:Chloroform (5:1, pH 4.5),

### Quantitative real-time PCR

Reverse transcription was performed using the Transcriptor First Strand cDNA Synthesis Kit (Roche) with oligo d(T) primers according to manufacturer’s protocol. The samples were incubated in a thermocycler at 25 °C for 10 min, then 55 °C for 30 min, and finally 5 min at 85 °C (C1000 Touch Thermal Cycler (Bio Rad)).

Real-time PCR was performed using the LightCycler 480 SYBR Green Master Kit (Roche). The sequences of the primers used are shown in Fig. S5B. The thermal profile was carried out according to default parameters: a pre-incubation step at 95 °C for 10 min (1 cycle), an amplification step (respectively at 95 °C for 10 s, 52 °C for 10 s, 72 °C for 20 s (45 cycles). All samples were analyzed in triplicate. UBC5 was used as reference genes. The amount of target was calculated using the following formula: amount of target = 2^−ΔΔCT^ (Livak and Schmittgen, 2001). For MeRIP, the amount of methylated transcripts level was determined by qRT-PCR using 2^-ΔCT^ methodology, the ΔCT value (cycle threshold) was normalized by ΔCT_IP_, ΔCT_input_ and input dilution factor. In order to determine the PCR efficiencies, standard curves for target and control genes were obtained using a series of cDNA dilutions as a template. The statistical significance of the results presented was estimated using a Student’s t-test at three significance levels: *p < 0.05, **p < 0.01, and ***p < 0.001.

### Library preparation and sequence analysis

Poly(A) RNA of the best quality from three replicates of each experiment were used for preparation of cDNA libraries for NGS using NEBNext Ultra II RNA Library Prep Kit for Illumina (NEB) according to the manufacturer’s protocol. Briefly we fragmented ≈ 5 ng of poly(A) RNA for 7 min, followed by cDNA synthesis, end repair, and adaptor ligation. The yield of amplified libraries was measured on a Qubit 4 fluorometer using the Qubit high sensitivity DNA kit (HS DNA kit) (Thermo Fisher Scientific). The libraries were sequenced at Genomed (Warsaw, Poland) using NovaSeq6000 (Illumina) in PE150 mode, 20 million paired reads per sample.

### Computational analysis of RNA-sequencing data

The quality of raw sequencing reads was assessed with FastQC v0.11.5. The quality trimming, filtering and adapter clipping were done with BBDUK2 v37.02 from BBMAP package. After discarding of rRNA-mapping reads using Bowtie 2 v2.3.5.1; the *L. angustifolius* ribosomal RNAs came from Ensembl Plants 56. The clean reads were then used for an ab initio transcriptome assembly with StringTie v1.3.3b and lupin genome LupAngTanjil_v1.0.

Expression levels of the assembled genes and transcripts were calculated with RSEM. Differential expression analysis and diagnostic plots were done using DESeq 2 in R/Bioconductor. Only genes displaying adjusted P-value < 0.05 were considered as differentially expressed. Additional plots were generated using own scripts in R, with ggplot and ggrepel libraries. Analysis of overrepresented Gene Ontology terms was done using clusterProfiler, separately for up- and downregulated genes. In each case, only 1000 genes with lowest adjusted P-value (as calculated with DESeq 2) and displaying at least two-fold change in expression were considered.

## Funding

This project was supported by Polish National Science Center (NCN) grant no. 2022/45/N/NZ9/04015

## Acknowledgments

The authors would like to thank C. Bousquet-Antonelli and R. Merret (Université de Perpignan Via Domitia, France) for their generous gift of anti-PAB2 antibodies and seeds of *ECT2-GFP* mutant of *Arabidopsis thaliana*, E. Gutierrez-Beltran for seeds of RBP47-GFP mutants of *Arabidopsis thaliana*, C. Schneider, Bruker Nano, (Fitchburg, Wisconsin, USA) and G. Szumilas, Labsoft (Poland) for the possibility to use the STORM system for these experiments.

## Notes

### Competing Interest Statement

The authors have declared no competing interest.

