## Supplemental for "Interplay between m6A modification and overall transcripts quantity: Impacts on mRNA composition in plant stress granules"

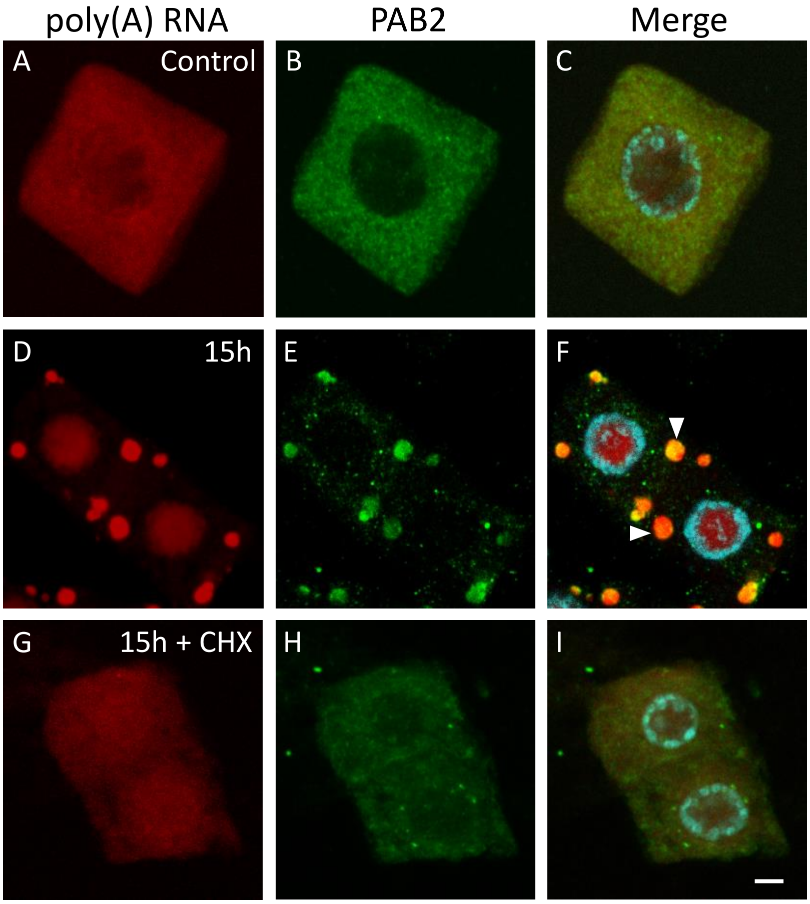


**Fig. S1** Distribution of poly(A) RNA (red fluorescence) and SGs marker protein PAB2 (green fluorescence) in meristematic cells of *L. angustifolius* roots in normoxia (**A-C**), 15 h of hypoxia (**D-F**) and 15 h hypoxia with cycloheximide treatment (**G-I**), the arrowheads indicate SGs, merge of signals and DAPI staining (**C**, **F**, **I**), bar 10 µm.

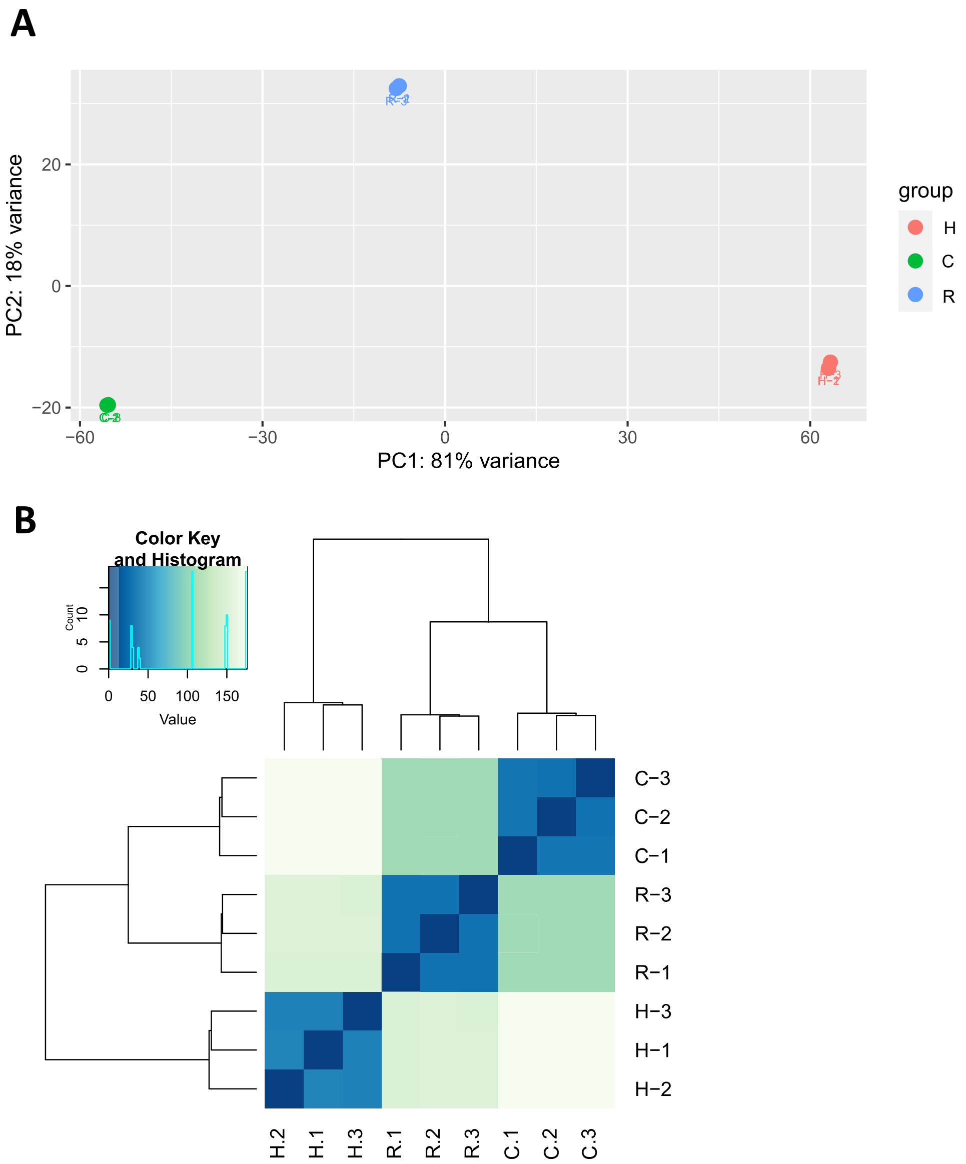


**Fig. S2** (**A**) Principal component analysis (PCA) of the RNA-seq data representing the clustering of biological replicates based on gene expression levels of *L. angustifolius* roots in normoxia (C), hypoxia (H) and reoxygenation (R) conditions. (**B**) Hierarchical gene clustering based on the Euclidean distance matrix between the replicates of above samples.

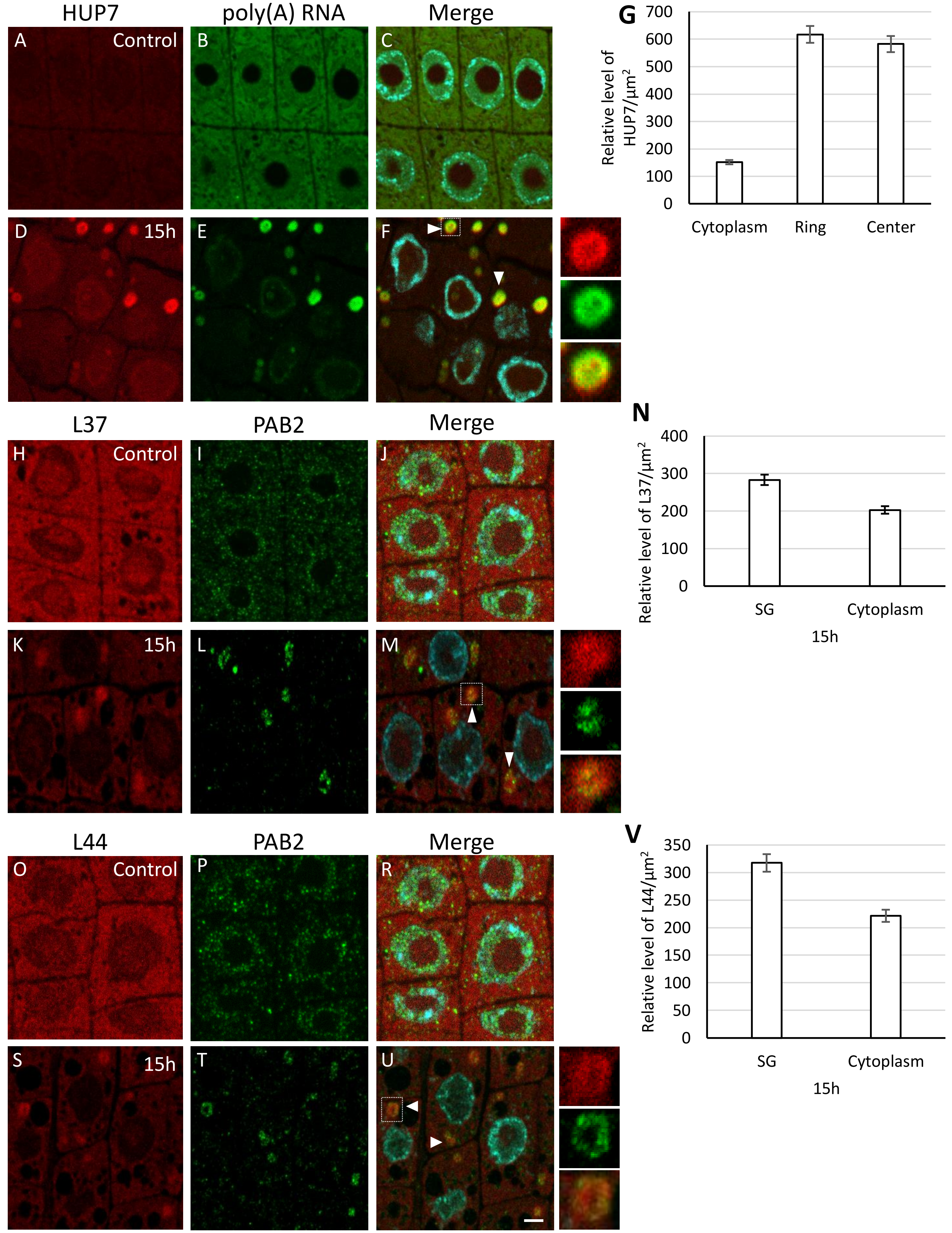


**Fig. S3** Localization of HUP7 mRNA (red fluorescence) and poly(A) RNA (green fluorescence) (**A-F**), L37 mRNA (red fluorescence) and SGs marker protein PAB2 (green fluorescence) (**H-M**), L44 mRNA (red fluorescence) and PAB2 (green fluorescence) (**O-U**) in meristematic cells of *L. angustifolius* roots in normoxia (**A-C**, **H-J**, **O-R**) and 15 h of hypoxia (**D-F**, **K-M**, **S-U**), the right-hand panel represents the magnification of SG which is marked with a square, merge of signals and DAPI staining (**C**, **F**, **J**, **M**, **R**, **U**), bar 10 µm. The relative fluorescence intensity of: HUP7 mRNA in the cytoplasm and ring and central area of SGs (**G**), L37 mRNA (**N**), L44 mRNA (**V**) in the cytoplasm and SGs of roots cells subjected 15 h to hypoxia stress.

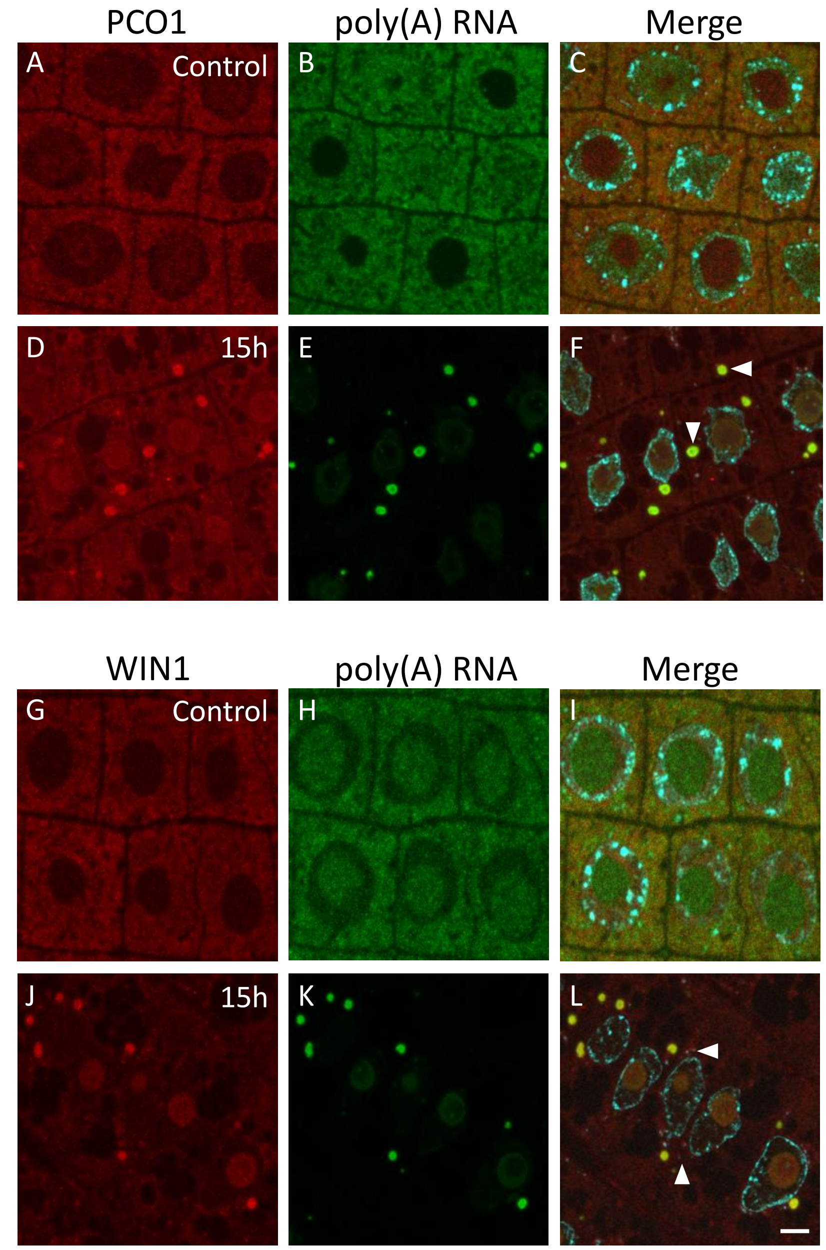


**Fig. S4** Detection of PCO1 transcripts (red fluorescence) and poly(A) RNA (green fluorescence) (**A-F**) and WIN1 mRNA (red fluorescence) and poly(A) RNA (green fluorescence) (**G-L**) in meristematic cells of *L. angustifolius* roots in normoxia (**A-C**, **G-I**) and 15 h hypoxia (**D-F**, **J-L**), the arrowheads indicate SGs, merge of signals and DAPI staining (**C**, **F**, **I**, **L**), bar 10 µm.

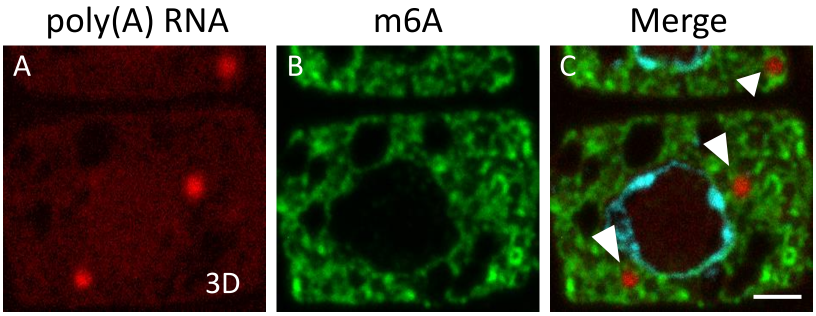


**Fig. S5** Localization of poly(A) RNA (red fluorescence, **A**) and m6A (green fluorescence, **B**) on resin sections in meristematic cells of *A. thaliana* 3 days hypoxia conditions, the arrowheads indicate SGs, merge of signals and DAPI staining (C), bar 10 µm.

**A**

| DNA probe | Sequence |
|---|---|
| ADH1-1 | CACCTGGTTTCAGATGAGTCACACCCTCTCCTA |
| ADH1-2 | GAAATCCTTGCCCCTTCAGCAGCAGCAAGGCC AACA |
| ADH1-3 | CCTATCAGTGTTGATCCTAAGAAGATC |
| HUP7 | TTGGCTAGTCGAGCTGTGTTGCCATGTGATAC AA |
| PCO1 | ATCAGCTAAAGTCCTCTCAGTACCCATTCAA |
| WIN | ACTGTGATGGTTGCTTGTTTTGTTGTTATCAGA |
| RPB1 | CATAATCATCTTTTGTAAAGTGAGGCAATGTCC GGTCTAT |
| L37 | TGATAATGAAGCCACTAAATATCATAGGAA |
| L44 | GTGCAGTGTATGCTTCTTGCATTCCTTGTTC |
| 18s rRNA | TTATCTAATAAATGCATCCTTCCAGGAAGTCG |
| 26s rRNA | TCCCGACAGGACGCTCTCACTCGAACCCTTC |

**Fig. S6** The sequences of antisense DNA probes (**A**) Sequence of primers used in qPCR reaction (**B**)

**B**

| Starter | Starter sequence 5'⟶ 3' |
|---|---|
| ADH1-1 | F: ATGAAGCTGGAGGGATTGTG<br>R: AGGTTCCGACAAAATGATGC |
| ADH1-2 | F: CATGAAGCTGGAGGGATTGT<br>R: CGAGGTTCCGACAAAATGAT |
| UBC5 | F:GAAATCGAGCGATGAAGAGC<br>R: CCCCTACCAGCAGCAATAAA |
| L37-1 | F: AGGGTTCTGCATCTGCATCT<br>R: ACGACTCTTCTGGAGGTGGA |
| L37-2 | F:ATGGGGAAGGGAACAGGTA<br>R: ACGAATTGCCTTCACACTCC |
| L44-1 | F: GCAAACAGTCCGGTTATGGT<br>R: CCCTTCTTGTCACCACCAAT |
| L44-2 | F: TGCAAGAACAAGGAATGCA<br>R: GCACTGCAACCTCAAGACAA |
| RPB1-1 | F: TTGGATTGAAACCCAGAAGC<br>R: TTCAGCCTCAAACACACTGC |
| RPB1-2 | F: TACCCCGAGACTGTGACTCC<br>R: ATGACGCTCCACCTTGTACC |
| HUP7-1 | F:CACGTCATTCCAAGAGCGAG<br>R:TCGTACATCAAATGCGCTGG |
| HUP7-2 | F:CCGCCGCTGAAATCTTGTAA<br>R:CCCCAAATGGCAAGGAAGAG |
| WIN-1 | F:CGTGCAGGGTCTAGTTCTGA<br>R:TGCTCCTCATCCAACCCATT |
| WIN-2 | F:GATGAGTGGAAGAAACGCCA<br>R:TCAGAACTAGACCCTGCACG |
| PCO1-1 | F:CGCTGACTCTTCCACCTTTG<br>R:GAGGACCAAGCACGTCTAGA |
| PCO1-2 | F:AACCGGTGGCGTACTAAGAA<br>R:ATACGGCATGTCAGGTGTCA |
